# FBXO11 deficiency in mice impairs lung development and aggravates cigarette smoke-induced airway fibrosis

**DOI:** 10.1101/2022.05.31.494154

**Authors:** Anitha Shenoy, Huacheng Luo, Sarah Lulu, Jennifer W. Li, Yue Jin, Qin Yu, Shu Guo, Vinayak Shenoy, Hao Chen, Andrew Bryant, Lizi Wu, Jia Chang, Kamal Mohammed, Jianrong Lu

## Abstract

Small airway fibrosis is a common pathology of chronic obstructive pulmonary disease (COPD) and contributes to airflow obstruction. However, the underlying fibrogenic mechanism is poorly understood. Epithelial-mesenchymal transition (EMT) has been proposed as a driver of fibrosis. EMT occurs in the airways of COPD patients and smokers, but it remains elusive whether EMT may contribute to airway fibrosis. We previously reported that FBXO11 is a critical suppressor of EMT and *Fbxo11* deficiency in mice causes neonatal lethality and EMT in epidermis. Here, we found that *Fbxo11*-deficient mouse embryonic lungs showed impaired epithelial differentiation, excess fibroblast cells surrounding the airways, and thickened interstitial mesenchyme. We further generated conditional mutant mice to ablate *Fbxo11* selectively in the club airway epithelial cells in adult mice, which induced partial EMT in the airways. To determine the effect of EMT on airway fibrosis, *Fbxo11* conditional mutant mice were exposed to cigarette smoke. Airway-specific loss of *Fbxo11* markedly enhanced smoking-induced airway fibrotic remodeling and collagen deposition. Taken together, our study suggests that EMT in the airway epithelium exacerbates cigarette smoke-induced airway fibrosis.

## Introduction

Tobacco smoke is a complex mixture of thousands of chemical compounds, many of which are cytotoxic, mutagenic, and proinflammatory. Of the numerous smoking-related maladies, the most common is chronic obstructive pulmonary disease (COPD). Chronic exposure to cigarette smoke is the primary risk factor for the development of COPD ^1,2^. COPD has become a global epidemic and is currently the third leading cause of death worldwide ^3^. COPD is a severely disabling lung disease characterized by persistent respiratory symptoms and airflow limitation that is due to airway and/or alveolar abnormalities ^4^. COPD typically involves pathological presentation in the airways (chronic bronchitis) and lung parenchyma (emphysema). Recent studies suggest that the narrowing of small conducting airways precedes the onset of emphysema and is an early prevalent clinical manifestation of COPD ^5-7^. Small-airway dysfunction is thus recognized as an important mechanism for COPD progression and the core pathology of COPD. Small airways in COPD patients commonly exhibit epithelial alterations, airway wall thickening, peribronchiolar fibrosis, persistent inflammation, and mucus hypersecretion. Fibrosis is a widespread feature of small airways in COPD patients and is considered the major contributor to small-airway narrowing in COPD ^5-7^, but the airway fibrogenic mechanism remains poorly understood.

Fibrosis is a chronic and progressive disorder defined by excessive deposition of extracellular matrix (ECM), which eventually leads to scar formation and organ failure ^8^. Myofibroblasts are generally thought of as the major type of ECM-producing cells in fibrosis ^9^. Epithelial-mesenchymal transition (EMT) has emerged as a crucial driver of tissue fibrosis ^10^. EMT is a dynamic cellular reprogramming process through which epithelial cells progressively shed their epithelial traits and shift toward a mesenchymal phenotype, including mesenchymal gene expression, elevated ECM production, and increased migratory capacity, invasiveness, and resistance to apoptosis. EMT is not a binary process. Activation of the EMT program is often partial, giving rise to cells with a continuous spectrum of hybrid epithelial-mesenchymal states ^11^. EMT is orchestrated by EMT-inducing transcription factors (EMT-TFs), mainly of the Snail, Zeb, and Twist families that are induced by an intricate network of pathways, including TGFβ signaling ^12^. EMT has been found to be associated with fibrosis in various organs. A prominent EMT inducer, TGFβ signaling is also a central mediator of the initiation and maintenance of fibrosis ^13^. The role of EMT in fibrogenesis was established by functional studies *in vivo*. Ectopic expression of *Snail* in the kidney or pancreas induces EMT and fibrosis in transgenic mice ^14,15^.

Conversely, genetic ablation of *Snail* or *Twist* significantly attenuates fibrosis in various mouse models ^16-18^. Mechanistically, type-2 EMT was initially thought to directly convert epithelial cells into fully mesenchymal myofibroblasts to drive fibrosis. However, this notion has largely been rebutted. Lineage tracing in genetically engineered mouse models indicates that epithelial cells with an EMT phenotype are not a significant source of myofibroblasts *in vivo* ^19-22^. Studies in mouse models of renal fibrosis suggest that injury induces tubular epithelial cells to undergo a partial EMT, which remain attached to the basement membrane, and secrete fibrogenic cytokines to promote inflammatory cell infiltration and renal fibroblast activation, leading to interstitial fibrosis ^17,18^.

The function of EMT in small-airway fibrosis of COPD remains to be investigated. Cigarette smoke condensate induces EMT in cultured human airway epithelial cells *in vitro* ^23-28^. Active EMT has been detected in the small and large airways of smokers and COPD patients ^25,27,29-31^. Moreover, genetic predisposition is an important risk factor for COPD ^32^. A human germline variant of *Snail* attenuates Snail’s ability to induce EMT and is associated with a decreased risk of COPD ^33^. However, there still lacks direct evidence supporting EMT’s contribution to airway fibrosis.

Animal models that recapitulate the main disease features are key to understanding the pathogenic mechanisms and discovering therapeutic targets. However, current laboratory animal models of COPD have been limited exclusively to the late emphysema and offer little relevance to the early airway disease of COPD ^2^. We and others previously reported that the F-box protein FBXO11, the substrate-recognition subunit of an E3 ubiquitin ligase complex, promotes the polyubiquitination and degradation of the Snail family members ^34,35^. FBXO11 is a critical suppressor of EMT and depletion of *Fbxo11* causes EMT in various epithelial cells *in vitro*. We genetically inactivated *Fbxo11* in mice, which led to neonatal lethality as well as aberrant Snail protein accumulation and partial EMT in the epidermis ^35^. In the present study, we showed that *Fbxo11*-deficient mouse embryos exhibited excess mesenchyme surrounding the airways and in the interstitium of the developing lung. To determine the role of EMT in the airway disease, we generated conditional mutant mice to specifically ablate *Fbxo11* in the airway epithelium, which were subsequently exposed to cigarette smoke. Airway-specific ablation of *Fbxo11* induced EMT and enhanced smoking-induced airway fibrosis in mice. The study thus provides functional evidence advocating EMT’s role in airway fibrosis and potentially a mouse model of airway disease.

## Materials and Methods

### Breeding of mice

*Fbxo11+/-* heterozygous mutant mice ^35^ were crossed with an FLP deleter strain 129S4/SvJaeSor-Gt(ROSA)26Sor^tm1(FLP1)Dym^/J (Jackson Laboratory, stock number: 003946) to generate *Fbxo11*^*fl*/+^ mice, which were bred to homozygosity (*Fbxo11*^*fl*/fl^) and with a club cell-specific Cre strain B6N.129S6(Cg)-Scgb1a1^tm1(cre/ERT)Blh^/J (Jackson Laboratory, stock number: 016225). Further breeding of Fbxo11^fl/fl^ mice with Fbxo11^fl/+^;Scgb1a1-Cre mice generated Fbxo11^fl/fl^;Scgb1a1-Cre “cKO” mice. To induce Cre, two-month-old mice were given tamoxifen (2 mg/mouse/day) by intraperitoneal injection for 5 consecutive days. PCR genotyping was performed using genomic DNA extracted from mouse lungs to verify the presence of the *Fbxo11* deletion allele.

### Exposure of mice to tobacco smoke

One week after tamoxifen injection, both *Fbxo11* cKO (Fbxo11^fl/fl^;Scgb1a1-Cre) mice and control littermates (Fbxo11^fl/fl^ or Fbxo11^fl/+^) were mock-treated or exposed to tobacco smoke as previously described ^38^, which included mainstream smoke and side stream smoke that were mixed with air, from 6 cigarettes (using 2R4F reference cigarettes from the Tobacco and Health Research University of Kentucky). Mainstream smoke is the smoke that comes out of un-lighted suctioned end of cigarette, while side stream smoke is the smoke that comes out of lighted end of the cigarette. The cigarette exposure in mice via nose closely mimics tobacco smoking in human. In order to expose cigarette smoke via nose only, we purchased a custom-made smoke exposure system (this system differed from conventional whole-body exposure systems). The manufacturer specially designed the mouse adaptor to minimize strain on animals. The adaptor holds a mouse in a resting position without movement. These adaptors are with 40 mm diameter and 110 mm long with a longitudinal slit and allow to hold mice with minimal strain.

Using the nose only inhalation system, the mice were allowed to inhale smoke from six cigarettes consecutively, 2 minutes for each cigarette with a 10 min interval for a total period of 62 minutes. During the period, mice remained in the adaptors. The smoke exposure was done for 5 days/week for up to 5 months. The body weights of all mice were measured weekly. After the mice were euthanized, the left lobe of the lungs was inflated with phosphate-buffered saline (PBS), followed by 4% Paraformaldehyde (PFA) and stored in 4% PFA overnight.

Subsequently, the fixed lung lobes were paraffin-embedded and sectioned. Parts of the right lobes were flash frozen for molecular assays. Mice were euthanized at various time points. For each time point, 3 mice from each genotype and treatment condition were analyzed.

### Collagen staining

Collagen staining was performed on 5 μm thick sections of 4% PFA fixed paraffin-embedded lung lobes using Picro-Sirius Red Stain Kit from ScyTek Laboratories Inc. Staining was performed using manufacturer’s protocol.

### Immunofluorescence

For immunostaining of lungs from adult mice, 4% PFA-fixed paraffin-embedded lung lobes were used. Sections (5 μm thick) were stained with anti-E-Cadherin (Cell Signaling, Cat No 3195), anti-Snail (Santa Cruz, Cat No Sc28199), and anti-Vimentin (Cell Signaling, Cat No 5741) after deparaffinization followed by citrate retrieval. For immunostaining of embryo lungs, fresh frozen OCT-embedded embryo lung sections (5 μm thick) were stained with anti-E-Cadherin (Cell Signaling, Cat No 3195), anti-Snail (Santa Cruz, Cat No Sc28199), and anti-Zeb1 (Santa Cruz, Sc 25388) after acetone retrieval. Images were taken using Leica DMI 6000 B microscope (Leica Microsystems) with the Openlab software (Perkin Elmer).

## Results

### FBXO11 deficiency in mouse embryos alters epithelial and mesenchymal development in the lung

We previously generated *Fbxo11* mutant mice in which a “STOP” cassette (consisting of a splice acceptor, LacZ-Neo, and the SV40 polyadenylation sequences) was inserted into the *Fbxo11* genomic locus, resulting in a null allele ^35^. *Fbxo11+/-* heterozygous mice appeared grossly normal and healthy. *Fbxo11-/-* homozygous mutant mice were born at the expected Mendelian ratios from heterozygous parents, but died shortly after birth.

Neonatal lethality is often owing to respiratory failure. Fetal lung development consists of several defined stages ^36^. During the saccular stage, the distal airways form saccular units. This stage is marked by progressive thinning of the interstitial mesenchyme and expansion of saccular airspace. Initially, the primary septa between the sacculi are thick. As the development proceeds, there is substantial interstitial thinning due to the disappearance of mesenchymal cells, and the sacculi turn into thin, smooth-walled sacs and are eventually remodeled into alveoli. The thinning of the interstitium juxtaposes the vascular capillaries with the epithelium. This process is critical for gas exchange at birth.

We characterized the developing lungs in the *Fbxo11* mutant embryos. Lung histology was performed on mouse embryos at E18.5 (saccular stage). Compared to wild-type control littermates, *Fbxo11-/-* homozygous mutants displayed an excessive number of subepithelial fibroblast-like cells surrounding the bronchi, resembling peri-bronchial fibrosis (Fig. 1A). Lungs from wild-type mice showed normal thinning of interstitial mesenchyme and formation of sac-like structures (pre-alveoli) (Fig. 1B). By contrast, lungs from *Fbxo11-/-* homozygous mutant embryos showed much thicker interstitial mesenchyme and less developed saccular structures (Fig. 1B). The thickened saccular walls resulted from an increased number of mesenchymal cells. The histological analysis suggests that global *Fbxo11* inactivation in mice leads to aberrant peri-bronchial and interstitial mesenchymal thickening in the lung at the saccular stage, which may cause respiratory failure and the observed neonatal mortality in the *Fbxo11-/-* mutant mice.

**Fig. 1.**
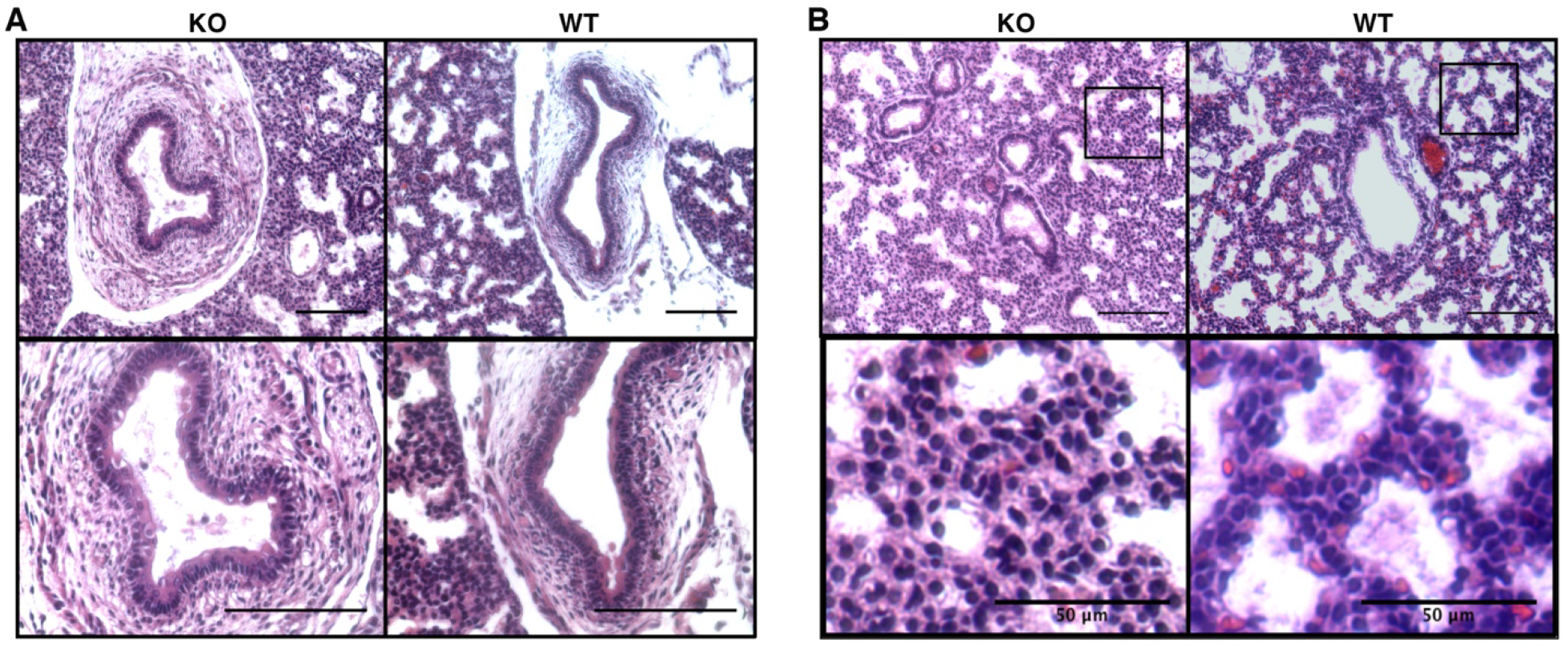
FBXO11 deficiency in mice causes thickening of subepithelial mesenchyme of conducting airways (**A**) and interstitial mesenchyme of saccular walls (**B**) in the developing lung. E18.5 embryonic lung sections from wild-type (WT) and *Fbxo11*-/-knockout (KO) littermates were stained by Hematoxylin & Eosin (H&E). The lower panels show higher magnification images. Scale bars: 100 μm (unless indicated).

Because of the excess mesenchyme phenotype in the *Fbxo11-/-* mutant lung, we performed immunofluorescence to compare the expression of epithelial and mesenchymal markers in E17.5 developing lungs of *Fbxo11+/-* heterozygous (control) and *Fbxo11-/-* homozygous littermates. The epithelial marker E-cadherin was strongly detected in airway epithelial cells in the control lung, but was largely diminished in *Fbxo11-/-* homozygous mutant lung (Fig. 2).

**Fig. 2.**
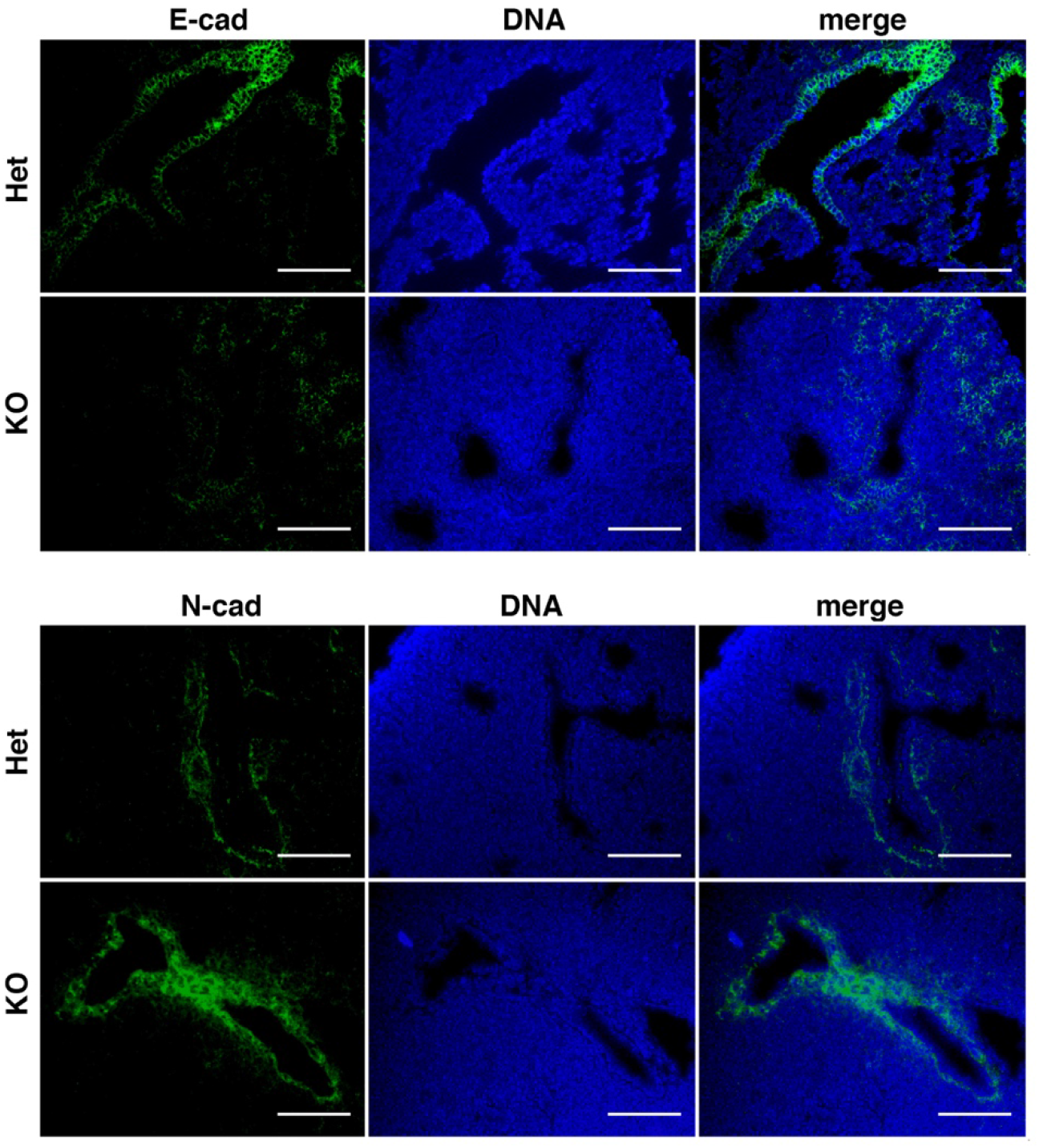
Altered epithelial and mesenchymal development in FBXO11-deficient mouse embryonic lungs. Sections through the lungs of E17.5 *Fbxo11*+/-(Het) control and *Fbxo11*-/-(KO) mouse embryos from the same litter were stained with antibodies against E-cadherin (E-cad) and N-cadherin (N-cad). Both cadherins show membrane-bound positivity in cells. DNA was stained by Hoechst 33342. Scale bars: 100 μm.

Expression of the mesenchymal marker N-cadherin was detected in cells beneath epithelium in the control lung, and was strongly enhanced in the lung of *Fbxo11-/-* homozygotes (Fig. 2). Taken together, the results suggest that FBXO11 deficiency in mice impairs airway epithelial differentiation and causes excess mesenchyme in the developing lung.

### Conditional deletion of FBXO11 in airway epithelium of adult mice leads to partial EMT

It is unclear if the epithelial and mesenchymal alterations in global *Fbxo11* inactivation mice are caused by cell-autonomous mechanisms. Therefore, we intended to ablate *Fbxo11* only in airway epithelium to determine if FBXO11 played a role in airway epithelial cells. The *Fbxo11* inactivation mutant allele had conditional potential. The STOP cassette was flanked by two FRT sites and could be removed by FLP recombinase-mediated recombination (Fig. S1). We bred *Fbxo11+/-* heterozygous mice with a FLP deleter strain that constitutively expressed FLPe (an enhanced variant of FLP) in most tissue types, including germ cells. We obtained offsprings carrying a conditional “floxed” (fl) *Fbxo11* allele in which exon 4 was flanked by LoxP sites and could be excised by Cre recombinase (Fig. S1). Homozygous *Fbxo11*^*fl/fl*^ mice were normal.

Exon 4 encodes for the F-box domain, which is essential for the assembly of the E3 ubiquitin ligase complex. Its deletion also causes a frameshift and loss of downstream protein sequence. Due to nonsense mutations, the exon 4-deleted mutant transcripts are likely subjected to nonsense-mediated decay. Collectively, deletion of exon 4 of *Fbxo11* generates a null allele.

In the mouse smaller conducting airways, the non-ciliated club cells (formerly known as Clara cells) are the predominant epithelial cell type lining the airway surface, which self-renew over the long term and give rise to ciliated cells ^37^. Therefore, we decided to ablate *Fbxo11* specifically in club cells so that the majority of airway epithelial cells would be *Fbxo11*-deficient. Club cells protect the bronchiolar epithelium by secreting surfactant proteins, including Secretoglobin 1A1 (SCGB1A1, also known as CCSP, CC10). Based on the lineage-restricted expression of Scgb1a1, Rawlins et al generated a “knock-in” transgenic mouse strain with a tamoxifen-inducible Cre recombinase (CreER) that was inserted into the *Scgb1a1* locus ^37^. Like the endogenous *Scgb1a1*, Scgb1a1-CreER was specifically expressed in the club epithelial cells in the airways and a small subset of type II cells in alveoli in the adult lung.

We bred *Fbxo11*-floxed mice with *Scgb1a1-CreER* mice to obtain *Fbxo11*^*fl/fl*^;*Scgb1a1-CreER* conditional knockout (cKO) mutant mice, which were born at the expected Mendelian ratio and appeared to be normal. When adult cKO mice (2-month old) were injected with tamoxifen to activate Cre in club cells, *Fbxo11* was ablated (based on PCR genotyping, Fig. S1). Due to a lack of FBXO11 antibodies suitable for immunostaining, we could not directly identify *Fbxo11*-null cells. Because FBXO11 is a part of ubiquitin ligase essential for Snail protein degradation, elevated Snail protein expression may serve as a surrogate indicator of FBXO11 deficiency. Two days after tamoxifen administration, we examined Snail and other molecular markers of EMT by immunostaining. Snail was weakly detected in the lung of littermate *Fbxo11*^*fl/*fl^ control mice, but displayed markedly increased expression in the cKO mice, mostly in airway epithelial cells (Fig. 3A), indicating successful deletion of *Fbxo11* in mouse airways. E-cadherin was robustly expressed in the airways of littermate control mice, but was largely diminished in the cKO mice (Fig. 3B). These cells with reduced E-cadherin expression remained in airway epithelium, suggesting that FBXO11 deficiency causes partial EMT. We further examined the mesenchymal marker Zeb1. In control mice, Zeb1 was barely detectable in the lung; In the cKO mice, strong Zeb1-positive mesenchymal cells were present surrounding airways (subepithelial) and in the interstitium (Fig. 3C). Because *Fbxo11* was deleted only in airway epithelium, the Zeb1+ mesenchymal cells apparently result from non-cell-autonomous function of FBXO11. Collectively, airway epithelial-specific ablation of *Fbxo11* induces partial EMT and affects surrounding mesenchyme.

**Fig. 3.**
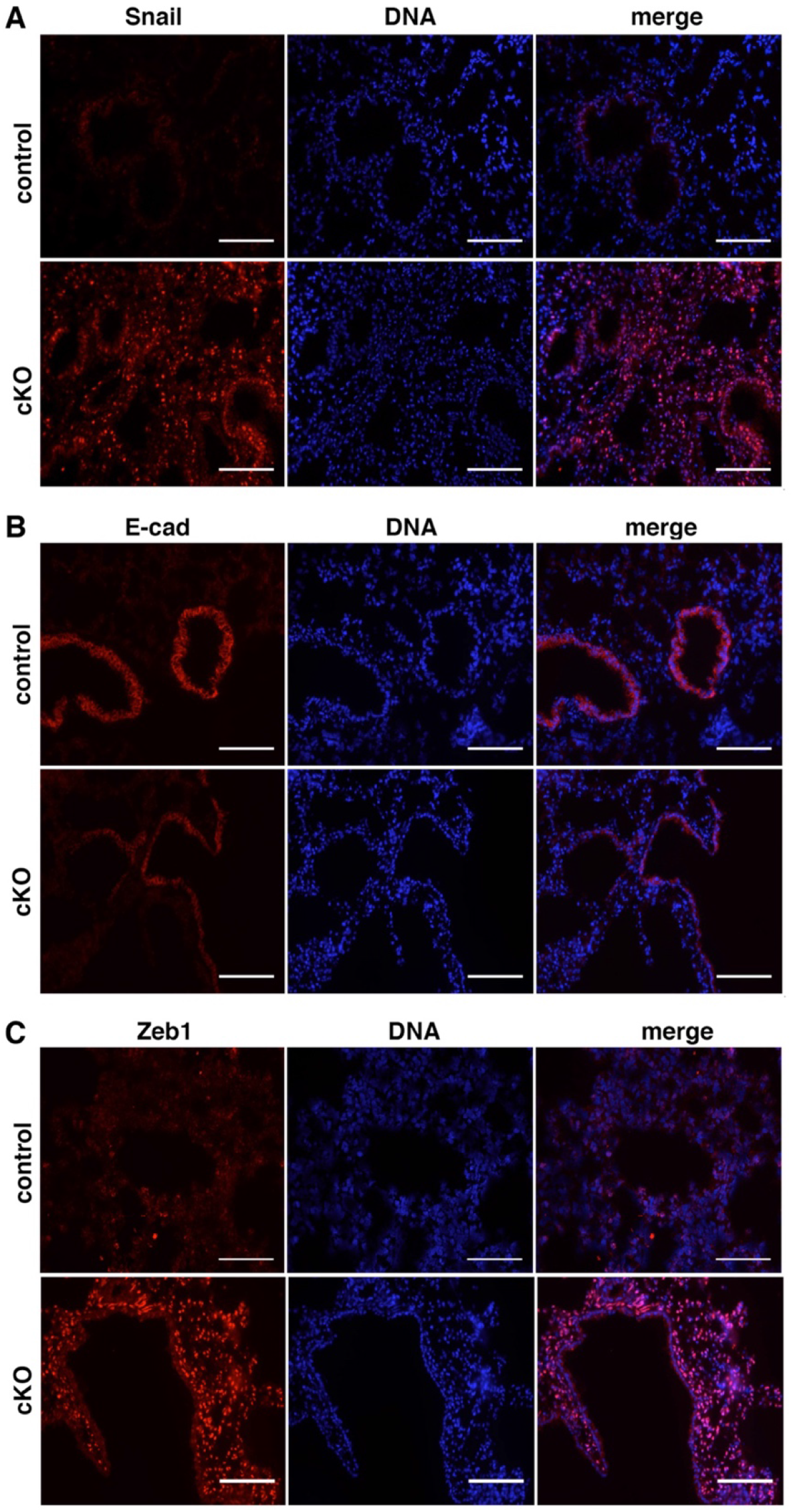
Ablation of *Fbxo11* specifically in airway epithelium of adult mice induces partial EMT. Control and *Fbxo11*^*fl/fl*^;*Scgb1a1-Cre* cKO mice were injected with tamoxifen for 5 days to activate Cre to delete *Fbxo11*. Two days after tamoxifen administration, mice were euthanized and lung sections were immuno-stained with indicated antibodies. Snail and Zeb1 show nuclear staining. Scale bars: 100 μm.

### FBXO11 deficiency enhances smoking-induced airway EMT and progressive fibrosis

The thickened mesenchyme surrounding the airways in the embryonic lung of global *Fbxo11* inactivation mutant mice was reminiscent of subepithelial fibrosis. Airway epithelial-specific deletion of *Fbxo11* in adult mice caused EMT. Therefore, we wondered whether EMT in airway epithelium due to FBXO11 deficiency might influence cigarette smoke-induced airway fibrogenesis. We decided to expose the *Fbxo11* airway cKO mutant mice to cigarette smoke and characterize the airway remodeling phenotype.

The mouse lung is a relatively stable organ with low rates of cell turnover, particularly in the airways. Following airway epithelial injury, cell loss is balanced with regeneration through amplification and differentiation of tissue stem and progenitor cells present within the adult lung. Club cells in small airways behave like progenitors for self-renewal in response to epithelial injury and differentiation into ciliated cells ^37^. Therefore, we expected that the *Fbxo11*-deficient club cells and derived ciliated cells would persist in the airways for a relatively long period. We established adult mouse cohorts with different genotypes: *Fbxo11*^*fl/fl*^;*Scgb1a1-CreER* cKO mice, and littermate control mice (*Fbxo11*^*fl/fl*^ or *Fbxo11*^*fl/+*^, without Cre). One week after tamoxifen administration, mice were maintained in regular room air or exposed to cigarette smoke (5 times each week) for up to 5 months. During the period, *Fbxo11* cKO mice did not show gross abnormalities. The cKO deletion allele was detected by PCR in genomic DNA extracted from mice at all experimental stages.

We first verified whether following cigarette smoke exposure, *Fbxo11* deficient cells in the cKO mouse lungs remained. Lungs from control and *Fbxo11* cKO mice that had been exposed to cigarette smoke for 2 months were analyzed by immunostaining. E-cadherin expression was particularly high in airway epithelium of control mice, but was markedly reduced in cKO lungs (Fig. S2). Meanwhile, mesenchymal markers Snail and N-cadherin exhibited markedly increased expression in cKO lungs compared with control lungs (Fig. S2). The result suggests that under cigarette smoke for 2 months, *Fbxo11*-deficient cells persist in the airways and exhibit a partial EMT phenotype. We then performed histological analysis of lungs from control and *Fbxo11* cKO mice without or with exposure to cigarette smoke for 2 months. Compared with control mice, *Fbxo11* cKO mice did not display clear histopathological changes in the airways (Fig. 4A). Smoke exposure did not cause evident morphological alterations in the lung of control mice, but slightly increased airway wall remodeling in the cKO mice (Fig. 4A). To evaluate potential fibrosis, we examined ECM deposition by collagen staining. Airways in the *Fbxo11* cKO mice showed moderately increased collagen deposition compared with control mice (Fig. 4A). Two months of smoke exposure caused more collagen deposition around the airways in the cKO mice than control mice (Fig. 4A). We further examined expression of α-smooth muscle actin (SMA). In all aforementioned lung samples, SMA appeared to be only expressed surrounding blood vessels (likely in pericytes) (Fig. 4B). The results suggest that 2-month smoke exposure leads to mildly increased epithelial remodeling and collagen accumulation in the airways of *Fbxo11* cKO mice.

**Fig. 4.**
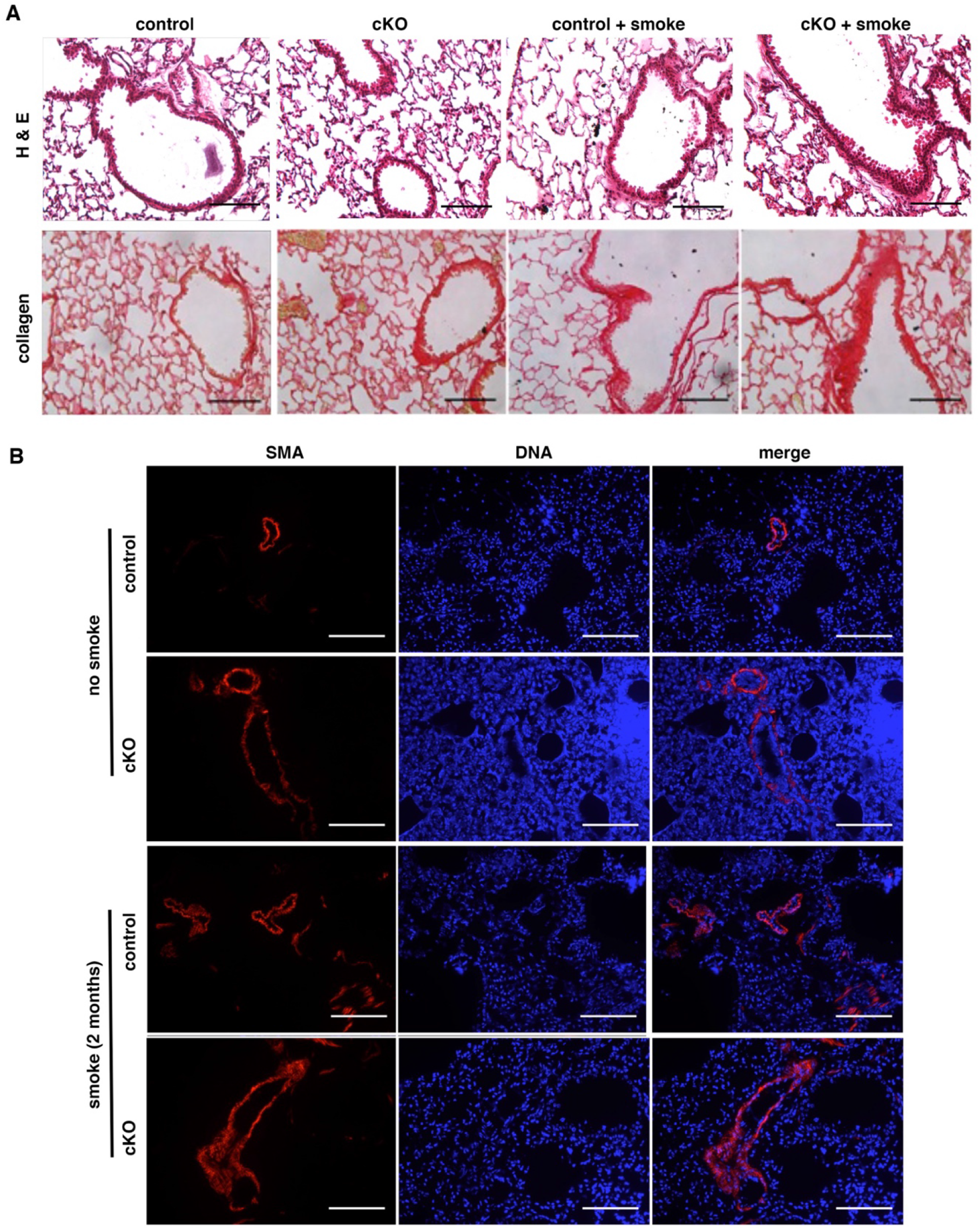
Airways in control and *Fbxo11* cKO mice following 2-month exposure to cigarette smoke. Control and *Fbxo11* cKO mice were exposed to cigarette smoke or regular air for 2 months. Lung sections were analyzed by H&E and collagen staining (**A**) and immuno-fluorescence of α-smooth muscle actin (SMA) (**B**). Scale bars: 100 μm.

Fibrosis is a progressive process. We thus characterized lung samples from mice with 5-month exposure to cigarette smoke. Immunostaining revealed that without smoke exposure, E-cadherin was strongly expressed in airways of control mice and was markedly downregulated by *Fbxo11* deletion in the cKO mice (Fig. 5A). Accordingly, Snail was markedly accumulated in the airways of cKO mice (Fig. 5A). Five-month exposure to cigarette smoke also downregulated E-cadherin in control mice, albeit without evident upregulation of Snail proteins (Fig. 5A), suggesting that smoke exposure may induce EMT-like changes in the airway. Interestingly, 5-month smoke exposure nearly eliminated E-cadherin expression in the airways of cKO mice (Fig. 5A), suggesting that prolonged cigarette smoke and *Fbxo11* deficiency each cause partial EMT in the airway epithelium, and their combination additively enhances the shift to a more complete EMT. Histologically, 5-month smoke exposure caused mild airway remodeling in the control mice, and more extensive airway wall thickening in the cKO mice (Fig. 5B). Five months of smoke exposure also mildly increased collagen deposition in the airways of control mice, but induced more fibrotic masses in the airways of *Fbxo11* cKO mice (Fig. 5B). Collectively, *Fbxo11* deficiency enhances the fibrogenic process induced by prolonged cigarette smoke exposure. The severity of fibrosis generally correlates with the extent of EMT.

**Fig. 5.**
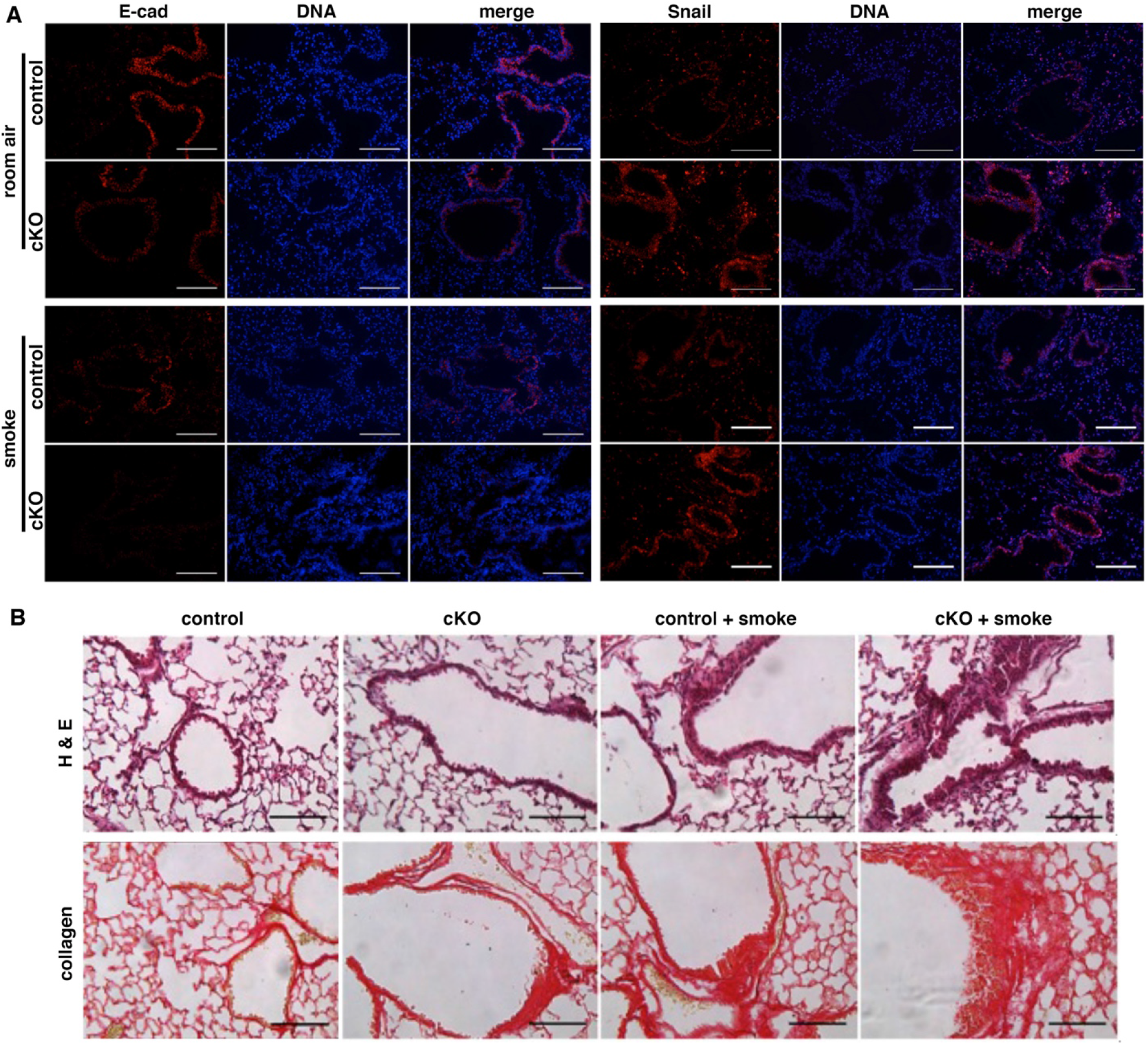
EMT in *Fbxo11*-deficient airway cells enhances cigarette smoke-induced fibrosis. Control and *Fbxo11* cKO mice were exposed to cigarette smoke or regular air for 5 months. Lung sections were immuno-stained for E-cadherin and Snail (**A**), or analyzed by histology and collagen staining (**B**). Scale bars: 100 μm.

## Discussion

Small airway fibrosis is an early and important pathological feature of COPD. Understanding and targeting airway pathogenesis may halt the progressive course of the disease. EMT is widely associated with tissue fibrosis and has been considered a key fibrogenic mechanism. EMT has been observed in the airways of COPD patients, but it remains elusive whether EMT indeed promotes airway fibrosis in COPD. In the present study, we genetically ablated *Fbxo11*, which encodes a critical suppressor of EMT, specifically in the mouse airway epithelium. The resulting *Fbxo11* mutant airways exhibited a partial EMT phenotype, validating FBXO11 as an essential suppressor of EMT in airway epithelium. Following prolonged exposure to cigarette smoke, such *Fbxo11* mutant mice with pre-existing EMT showed a much stronger airway fibrotic response than their regular littermates. Therefore, the present study provides *in vivo* evidence that EMT can enhance smoking-induced pathogenesis of airway fibrosis. EMT may represent a potential therapeutic target for the treatment of airway fibrosis in COPD.

Although cigarette smoking is the major etiological factor for COPD, only a small subset of smokers develop COPD ^2^. The cause of individual susceptibility is not fully elucidated ^2^. Genetic predisposition is also a risk factor. As EMT contributes to the airway disease of COPD, individuals with an increased tendency to undergo EMT may be predisposed to smoking-induced airway fibrogenesis. Consistent with this notion, a human genetic variant of Snail is associated with COPD vulnerability ^33^.

Animal models are a valuable tool to elucidate the pathogenic mechanisms. However, there lack experimental mouse models that resemble airway pathogenesis of COPD. The airway-specific *Fbxo11* deletion mutant mice may serve as a sensitized animal model to study cigarette smoke-induced airway disease. However, even in the *Fbxo11* cKO mutant mice following 5-month smoking, the airway fibrotic phenotype is still relatively moderate and may mimic the early stage of fibrosis. Chronic exposure to cigarette smoke (e.g., more than one year) may be necessary to reproduce the full-blown airway fibrosis in COPD patients.

## Acknowledgments

The work was in part supported by Florida James and Esther King Biomedical Research Program (4KB07 to JL).

